# Wall teichoic acid substitution with glucose governs phage susceptibility of *Staphylococcus epidermidis*

**DOI:** 10.1101/2023.07.27.550822

**Authors:** Christian Beck, Janes Krusche, Anna Notaro, Axel Walter, Lara Kränkel, Anneli Vollert, Regine Stemmler, Johannes Wittmann, Martin Schaller, Christoph Slavetinsky, Christoph Mayer, Cristina De Castro, Andreas Peschel

**Author notes:** Correspondence to Andreas Peschel.

## Abstract

The species- and clone-specific susceptibility of *Staphylococcus* cells for bacteriophages is governed by the structures and glycosylation patterns of wall teichoic acid (WTA) glycopolymers. The glycocodes of phage-WTA interaction in the opportunistic pathogen *Staphylococcus epidermidis* and in other coagulase-negative staphylococci (CoNS) have remained unknown. We report a new *S. epidermidis* WTA glycosyltransferase TagE whose deletion confers resistance to siphoviruses such as ΦE72 but enables binding of otherwise unbound podoviruses. *S. epidermidis* glycerolphosphate WTA was found to be modified with glucose in a *tagE*-dependent manner. TagE is encoded together with the enzymes PgcA and GtaB providing uridine diphosphate-activated glucose. ΦE72 transduced several other CoNS species encoding TagE homologs suggesting that WTA glycosylation via TagE is a frequent trait among CoNS that permits inter-species horizontal gene transfer. Our study unravels a crucial mechanism of phage-*Staphylococcus* interaction and of horizontal gene transfer and it will help in the design of anti-staphylococcal phage therapies.

**Importance:** Phages are highly specific for certain bacterial hosts, and some can transduce DNA even across species boundaries. How phages recognize cognate host cells remains incompletely understood. Phages infecting members of the genus *Staphylococcus* bind to wall teichoic acid (WTA) glycopolymers with highly variable structures and glycosylation patterns. How WTA is glycosylated in the opportunistic pathogen *Staphylococcus epidermidis* and in other coagulase-negative *Staphylococcus* (CoNS) species has remained unknown. We describe that *S. epidermidis* glycosylates its WTA backbone with glucose and we identify a cluster of three genes, responsible for glucose activation and transfer to WTA. Their inactivation strongly alters phage susceptibility patterns, yielding resistance to siphoviruses but susceptibility to podoviruses. Many different CoNS species with related glycosylation genes can exchange DNA via siphovirus ΦE72 suggesting that glucose-modified WTA is crucial for interspecies horizontal gene transfer. Our finding will help to develop antibacterial phage therapies and unravel routes of genetic exchange.

## Introduction

*Staphylococcus epidermidis* is one of the most abundant colonizers of mammalian skin and of nasal epithelia [1, 2]. Some nosocomial *S. epidermidis* clones also cause invasive infections, in particular biofilm-associated infections on catheters or artificial implants such as hip and knee joints or heart valves [3, 4]. Although *S. epidermidis* is not an as aggressive pathogen as *Staphylococcus aureus*, biofilm-associated infections are difficult to treat and cause a high burden of morbidity and costs for health care systems. Many *S. epidermidis* clones are resistant to beta-lactams and other antibiotics such as linezolid, which further complicates the treatment of *S. epidermidis* infections [1].

The major invasive *S. epidermidis* clones seem to pursue two different virulence strategies. The MLST type 2 (ST2) strains produce particularly strong biofilms [3, 5]. In contrast, ST10, ST23, and ST87 clones are only weak biofilm formers, but they express an additional surface molecule that alters their host interaction capacities and leads to a shift from commensal to pathogen behavior [6]. Surface properties and host interaction of staphylococci are governed not only by surface proteins but also by cell-wall anchored glycopolymers composed of alditolphosphate repeating units called wall teichoic acids (WTA) [7, 8]. The WTA polymers of *S. epidermidis* and other coagulase-negative *Staphylococcus* (CoNS) species have remained a neglected field of research despite their potentially critical role for host colonization and infection. Most *S. epidermidis* clones seem to express WTA composed of glycerolphosphate (GroP) repeating units [9]. A recent study has shown that ST10, ST23, and ST87 strains express an additional *S. aureus*-type WTA composed of ribitolphosphate (RboP) repeating units, which shapes their interaction with human epithelial and endothelial cells [6].

WTA is also crucial for binding of virtually all known *Staphylococcus* phages, which use differences in WTA structure to recognize their cognate host species [10]. Phages of the Sipho- and podovirus groups often not only discriminate between different WTA backbones but also between different types of backbone glycosylation. Most Firmicutes link D-alanine esters and sugar residues to GroP or RboP repeating units [7, 8]. Variation in glycosylation for instance by N-acetylglucosamine (GlcNAc) in alpha or beta configuration or N-acetylgalactosamine (GalNAc) has been found to govern the susceptibility patterns of *S. aureus* strains for different phages [11–14]. The group of broad-host range myoviruses, however, requires WTA for binding but does not discriminate between RboP and GroP WTA and does not require WTA glycosylation [15–17].

WTA-phage interaction is of importance for phage-therapeutic strategies, which have gained increasing attention recently [3, 18]. Moreover, they are critical for inter-species horizontal gene transfer via transducing bacteriophages [19]. Such transduction events have led to the transfer of resistance and virulence genes into the genomes of *S. aureus* and other species, thereby allowing, for instance, evolution of methicillin-resistant *S. epidermidis* (MRSE) and methicillin-resistant *S. aureus* (MRSA) [20, 21]. Despite the critical role of WTA in these processes, the biosynthesis, composition, and glycosylation of the canonical *S. epidermidis* WTA has not been studied.

Here we demonstrate, that *S. epidermidis* strain 1457 glycosylates its GroP-WTA with glucose and we identify the WTA glucosyltransferase gene *tagE*. *S. epidermidis tagE* mutants showed complex changes in phage susceptibility patterns including both, the loss, and the acquisition of susceptibility to certain phages, some of which we found to be capable of transducing plasmid DNA between different CoNS species.

## Results

### 1. Disruption of a putative glycosyl transferase gene cluster confers resistance to phage ΦE72

Several new phages with the capacity to infect *S. epidermidis* have been reported recently [22, 23]. Some of them have the capacity to transduce DNA between different *S. epidermidis* lineages, raising the question, which bacterial target structures are recognized by the phages’ binding proteins, and how universal these target structures may be among different clones of *S. epidermidis* and other CoNS. As most *S. aureus* phages recognize the sugar modifications of WTA [11–13, 24], it was tempting to speculate that glycosylated GroP-WTA is also required for binding of *S. epidermidis* phages. However, the enzymes responsible for WTA glycosylation in *S. epidermidis* have remained unknown and it has also remained elusive, which glycosylation patterns can be found on *S. epidermidis* GroP-WTA. To elucidate the WTA glycosylation pathways of *S. epidermidis* and explore its impact on phage interaction we set out to identify and inactivate the responsible enzyme genes.

A library of transposon mutants of *S. epidermidis* 1457 was created using a xylose-inducible Himar1 transposase [25] and incubated with phage ΦE72, which is known to infect and multiply in strain 1457 [22]. Two mutants, which were resistant to ΦE72 were identified and found to have the transposon integrated in two adjacent genes of unknown function (Fig. 1a; Fig. 2a, d). The two genes were not in the vicinity of other WTA-biosynthesis related genes, but their gene products shared similarity with glycosylation-related enzymes. The gene B4U56_RS02220 product was 46% similar to TagE of *Bacillus subtilis*, which glycosylates GroP-WTA with glucose residues [26] and 48% similar to TarM of *S. aureus,* which glycosylates RboP-WTA with GlcNAc (Fig. 1b) [27]. The adjacent gene B4U56_RS02215 encodes a protein with 59% similarity to the phosphoglucomutase PgcA of *B. subtilis*, which isomerizes glucose-6-phosphate to yield glucose-1-phosphate [28]. In addition, the product of gene B4U56_RS02210, next to *pgcA,* was 85% similar to the GtaB enzyme of *B. subtilis* generating UDP-glucose from glucose-1-phosphate and UTP [29]. Both, PgcA and GtaB are required for glycosylation of GroP-WTA with glucose via TagE in *B. subtilis* [30], although the two genes are not encoded together with *tagE* in *B. subtilis* [31, 32]. We assumed that the three enzymes might cooperate in *S. epidermidis* to activate and attach glucose to GroP-WTA.

**Figure 1:**
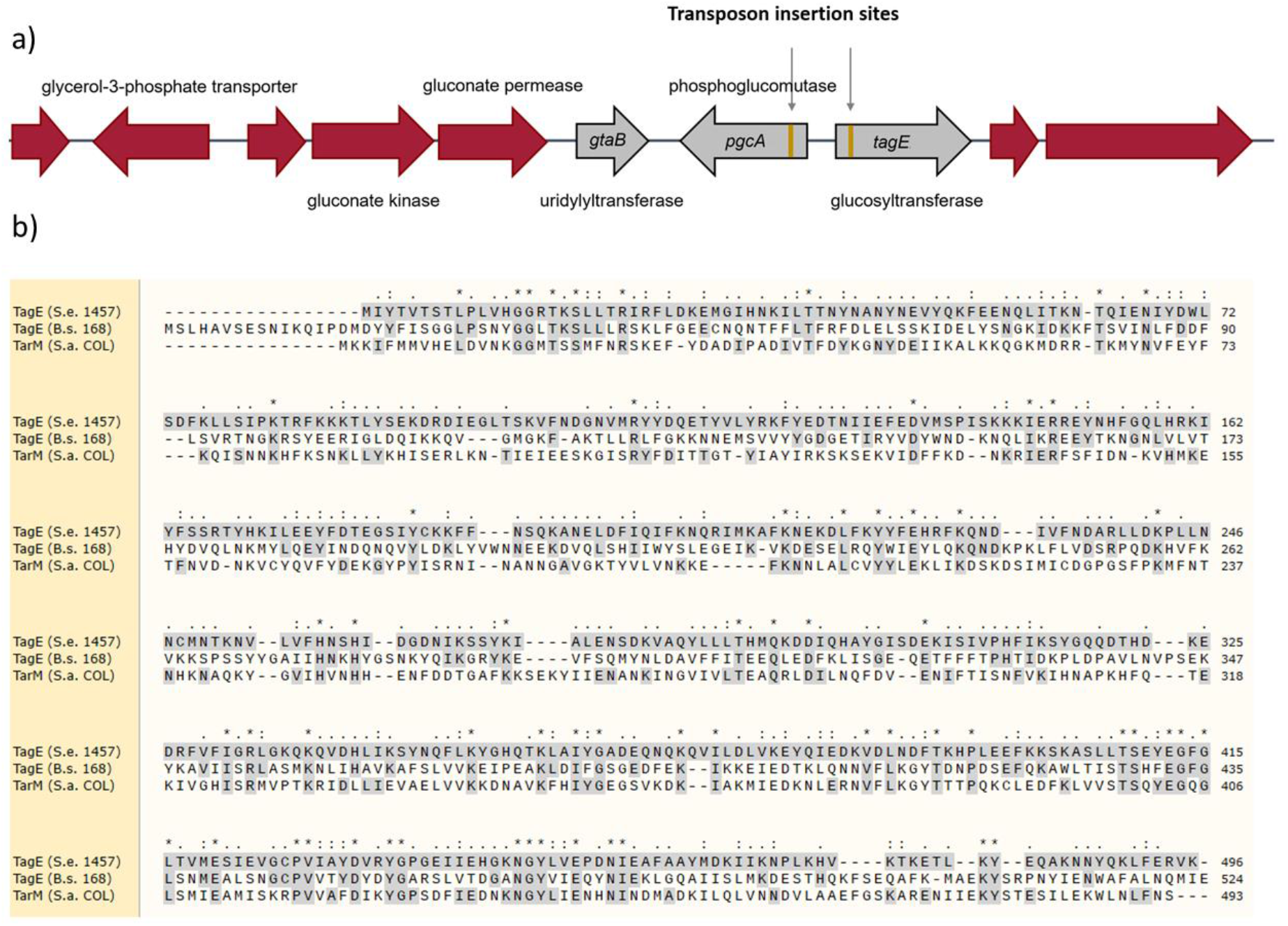
The *tagE* gene encodes a glycosyltransferase in *S. epidermidis*. a) Genetic locus identified by transposon mutagenesis contains the *S. epidermidis tagE, pgcA,* and *gtaB* homologues. Transposon insertion sites are labeled in gold. b) MUSCLE alignment of *S. epidermidis* TagE with *B. subtilis* TagE and

**Figure 2:**
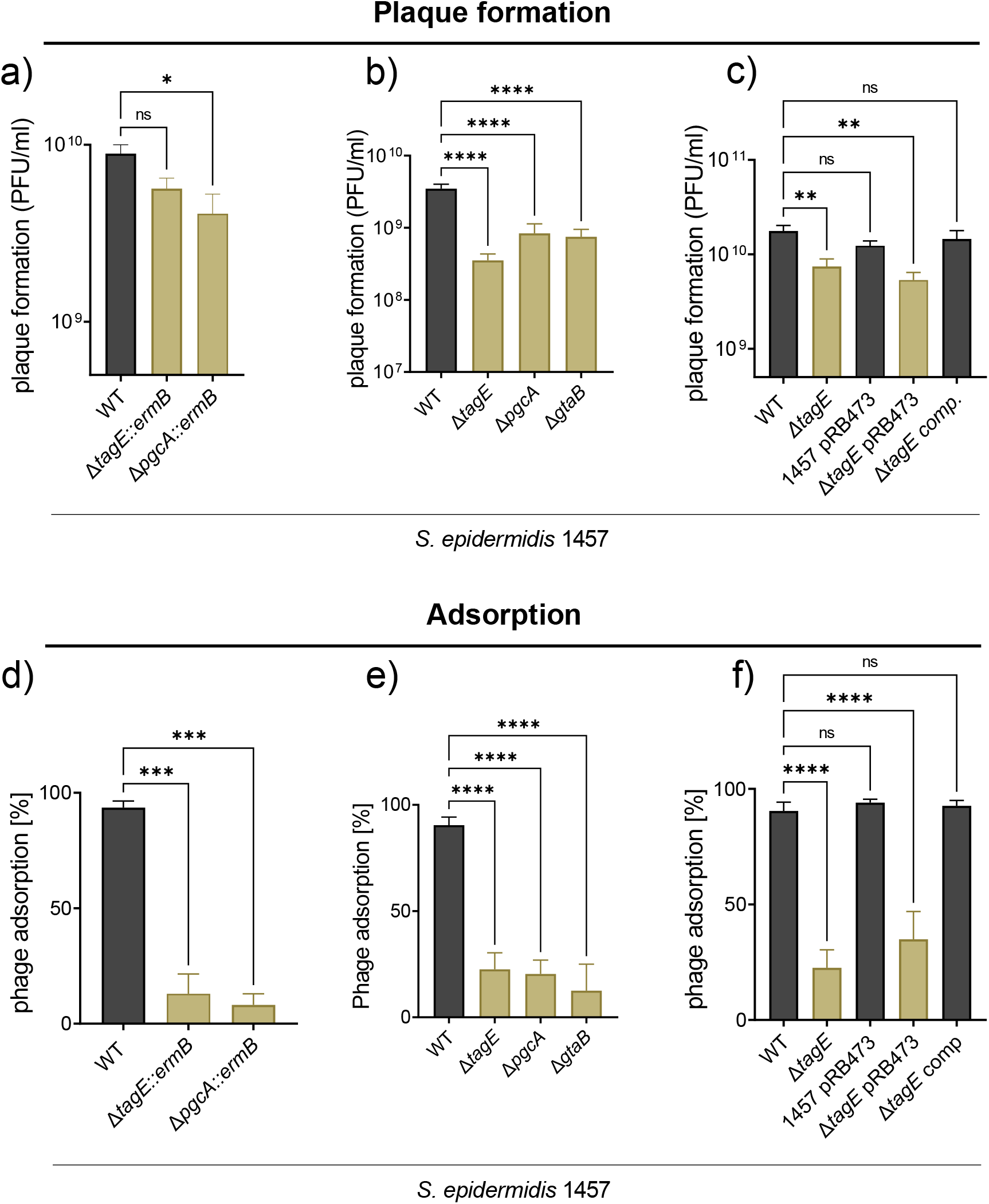
ΦE72 shows decreased infection (a,b) and binding (d,e) of the *tagE*, *pgcA,* and *gtaB* mutants. This deffect can be restored by complementing the *tagE* mutant with the genetic locus containing *tagE*, *pgcA,* and *gtaB* on plasmid pRB473 (c,f). The data represent the mean ± SEM of at least three independent experiments. Ordinary one-way ANOVA was used to determine statistical significance versus *S. epidermidis* 1457 wild type (WT), indicated as: not significant (ns), *P < 0.05, **P < 0.01, ***P < 0.001, ****P < 0.0001.

### 2. *S. epidermidis* TagE is responsible for glucose addition to GroP-WTA

The three *S. epidermidis* genes were renamed according to the corresponding *B. subtilis* genes *tagE*, *gtaB*, and *pgcA.* All three genes were inactivated by targeted deletion to confirm their roles in phage susceptibility. The three mutants were as resistant to ΦE72 infection as the transposon mutants, and complementation of the *tagE* mutant with a plasmid-encoded copy of the gene locus restored wild-type level ΦE72 susceptibility (Fig. 2 c,f). The various transposon and targeted deletion mutants were approximately 3-fold less susceptible to ΦE72 infection, but were not completely resistant, suggesting that the phage may have additional, albeit less effective ways to interact with *S. epidermidis* 1457. In a similar way, and even more pronounced, the mutants had retained only limited capacities to bind ΦE72 particles in liquid media (Fig. 2 d-f; Fig S1).

WTA isolated from 1457 wild type (WT) contained substantial amounts of glucose when analyzed by an enzymatic glucose assay indicating that ca. 50% of the GroP-WTA repeating units are modified with glucose (Fig. 3a). In contrast, none of the WTA samples of any of the *tagE*, *gtaB*, or *pgcA* mutants was found to contain glucose. High-performance liquid chromatography coupled to a mass spectrometry detector (HPLC-MS) and nuclear magnetic resonance (NMR) spectroscopy confirmed the presence of glucose-substituted GroP repeating units in the wild type and the absence of glucose in the mutants (Fig. 3b,c; Fig. S2). These findings reflect earlier reports on the presence of glucose on *S. epidermidis* GroP-WTA [9] and they confirm that the PgcA-GtaB-TagE pathway is required for GroP-WTA glycosylation with glucose. NMR analysis indicated that the glucose units are α configured at the anomeric center and attached to the C2-position of GroP. About 15% of the glucose residues are modified with D-alanine at the O6-position of glucose (Fig. 3c; NMR extended description). The α-configuration is reminiscent of the configuration of GlcNAc on RboP-WTA introduced by the TagE-related TarM in *S. aureus* [27].

**Figure 3:**
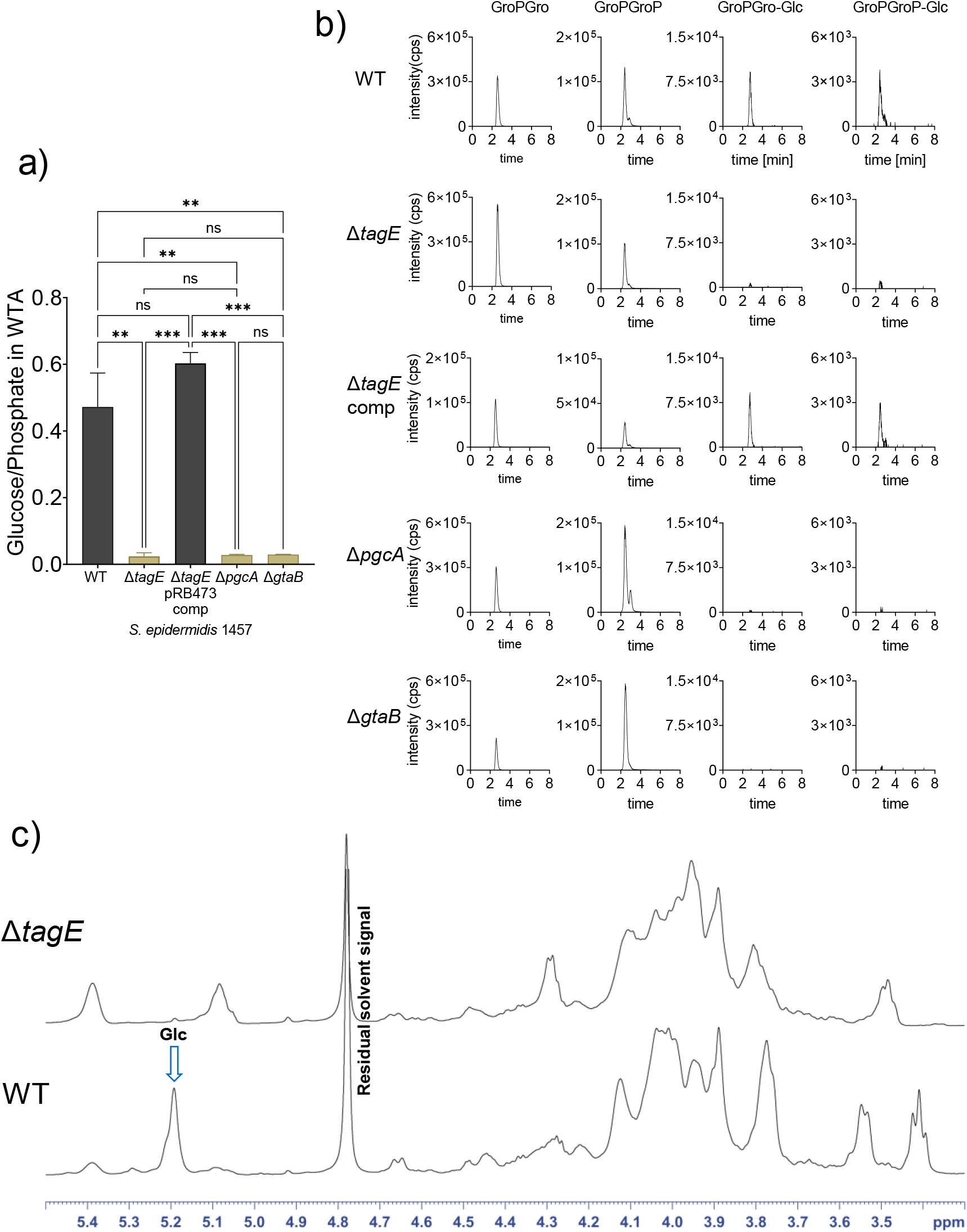
WTA analysis of the *S. epidermidis* mutants Δ*tagE*, Δ*pgcA,* Δ*gtaB* and of Δ*tagE* containing the pRB473 plasmid carrying *tagE, pgcA, and gtaB* genes for complementation. a) Ratio of glucose per phosphate content of WTA measured enzymatically. b) HPLC-MS: Extracted ion chromatograms (EIC) of GroP-Gro ([M - H]^−^ = 245.0432) and GroP-GroP ([M - H]^−^ = 325.0095) with (GroP-Gro-Glc; [M - H]^−^ = 407.096) (GroP-GroP-Glc; [M - H]^−^ = 487.0623) or without glucose substitution. c) ^1^H NMR spectra reveal D-glucose on WTA of the *S. epidermidis* 1457 wild type (WT) (at the C2-position of GroP), while deletion of *tagE* results in absence of glucose on WTA. For a) data represent the mean ± SEM of at least three independent experiments. Ordinary one-way ANOVA was used to determine statistical significance, indicated as: not significant (ns), **P < 0.01, ***P < 0.001.

The absence of glucose on GroP-WTA in the Δ*tagE* mutant did not alter biofilm formation by *S. epidermidis* 1457 (Fig. S3). Moreover, no differences in growth kinetics (Fig. S1), cell wall thickness, or cell shape (Fig. S4) were observed in the mutants, indicating that the absence of glucose on GroP-WTA has no major impact on overall cellular properties of the *S. epidermidis* surface.

UDP-glucose generated by PgcA and GtaB is also required for biosynthesis of the glycolipid diglucosyldiacylglycerol (DGlcDAG), which serves as anchor structure for lipoteichoic acid (LTA) polymers in *B. subtilis* and many other Firmicutes (Fig 4a) [33–35]. However, DGlcDAG is not essential for LTA biosynthesis because mutants lacking the glycolipids still produce LTA attached to phosphatidylglycerol lipids [35, 36]. The *S. epidermidis pgcA* and *gtaB* mutants, but not the *tagE* mutant, also lacked DGlcDAG, which was present in the parental strain (Fig. 4b), indicating that DGlcDAG is synthesized in *S. epidermidis* by the same pathway as in *B. subtilis* and *S. aureus*.

**Figure 4:**
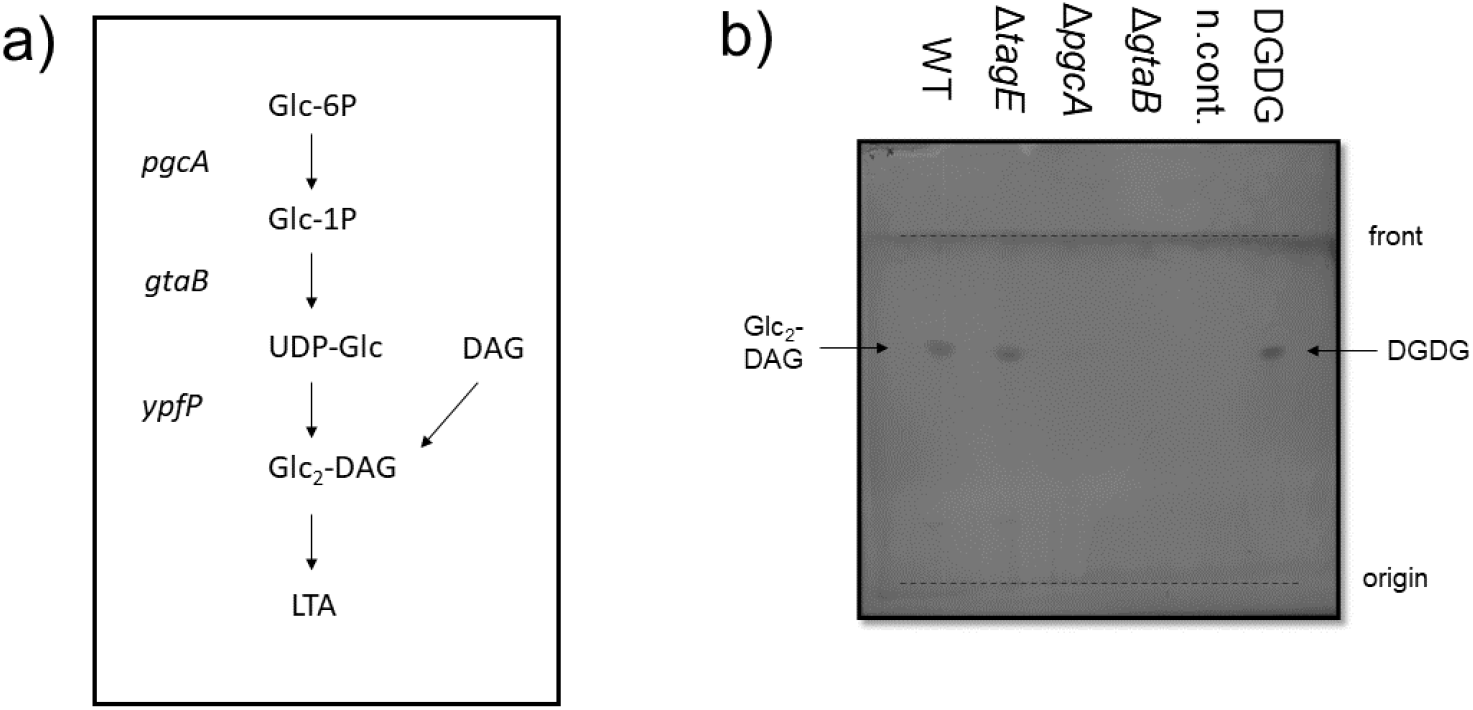
Glycolipid detection by TLC. a) LTA glycolipid biosynthesis pathway as described for *S. aureus* and *B. subtilis* (adapted from [36]). b) Glycolipid detection on a TLC plate stained with α-naphtol/sulfuric acid. 5 µg of digalactosyldiacylglycerol (DGDG) was used as positive control, the solvent methanol/chloroform (1:1) as negative control (n.cont.). One representative experiment of three independent experiments is shown.

### 3. Lack of WTA glucose impairs binding of known *S. epidermidis* siphoviruses but promotes binding of podoviruses

Several other phages in addition to ΦE72 were analyzed for an impact of GroP-WTA glucose modification on phage binding and infection. The ΦE72-related siphoviruses Φ456, Φ459, and Φ27, which are known to bind to *S. epidermidis* 1457 [22], showed reduced binding to the *pgcA, gtaB*, and *tagE* mutants compared to the wild type but the reduction was less pronounced as for ΦE72 (Fig. 5a,f). Φ459 was equally reduced in its capacities to propagate in the mutants as ΦE72 (Fig. 5b). Despite their capacity to bind *S. epidermidis* 1457, Φ27 and Φ456 did not form clear plaques on wild-type or mutant strains. Two recently isolated myoviruses of the genus sepunavirus, ΦBE04 and ΦBE06 [37], showed no reduction in their ability to bind and infect the mutants, suggesting that these myoviruses are not dependent on glucose-modified GroP-WTA (Fig. 5d,e). This behavior resembles the lacking impact of WTA glycosylation on myovirus ΦK infection of *S. aureus* [13].

**Figure 5:**
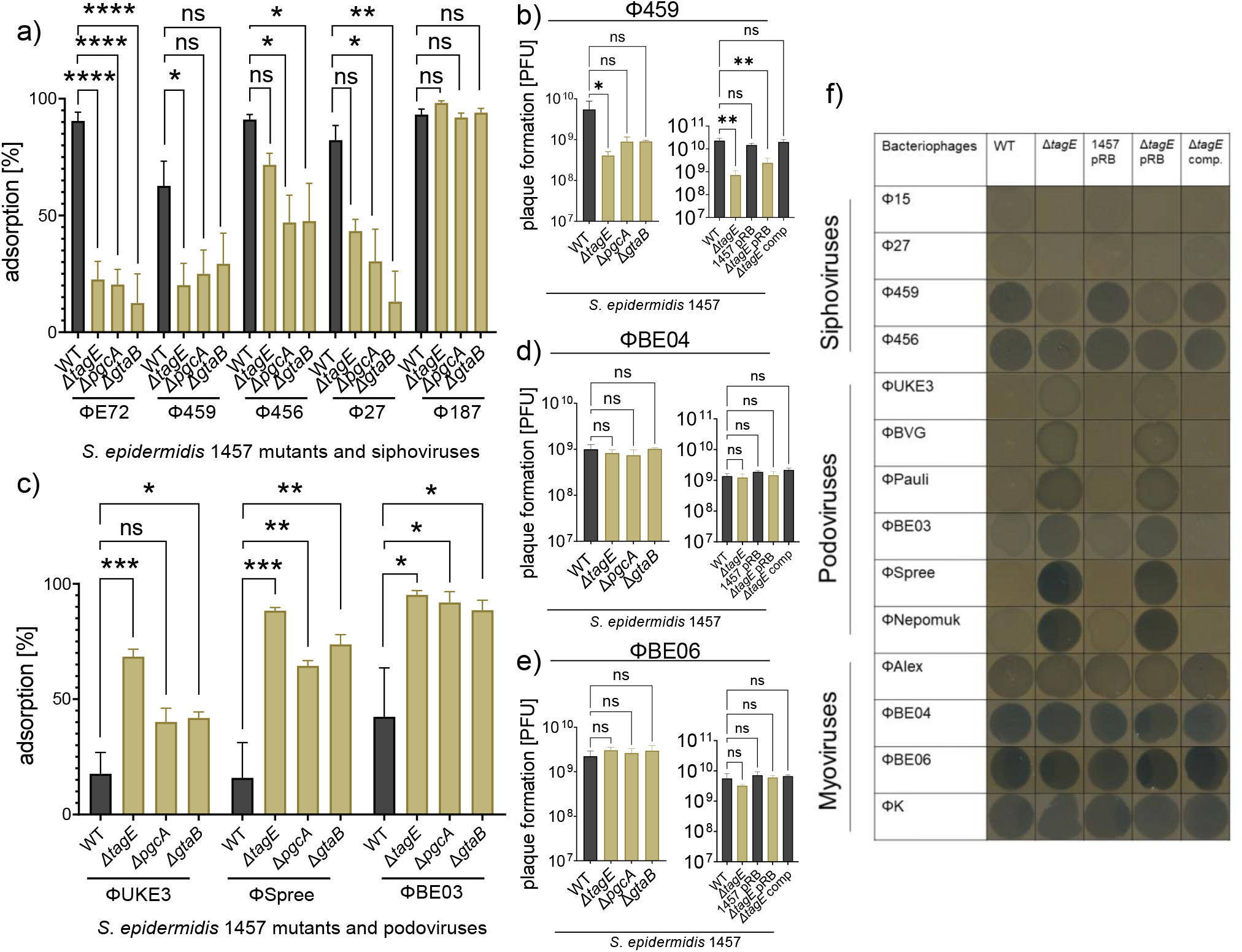
TagE-glycosylated WTA increases binding of siphoviruses but reduces podovirus binding. WTA glycosylation-deficient mutants of *S. epidermidis* show decreased binding of ΦE72-related siphoviruses Φ459, Φ456, and Φ27 (a), but increased binding of the podoviruses ΦUKE3, ΦSpree, and ΦBE03 (c), while the GroP-GalNAc-specific siphovirus Φ187 still shows strong binding (a). WTA glycosylation-deficient mutants of *S. epidermidis* show less plaque formation by ΦE72-related siphovirus Φ459 (b), while plaque formation by the myoviruses ΦBE04 (d) and ΦBE06 (e) remains unchanged. f) Lytic zones and “lysis from without” by siphoviruses decrease in the absence of *tagE* but increase for podoviruses. Myoviruses show formation of lytic zones independently of the presence or absence of *tagE.* (pRB= pRB473 (empty vector control); comp = complementation with *tagE*, *gtaB*, *pgcA* genes) The data represents the mean ± SEM of at least three independent experiments. Ordinary one-way ANOVA was used to determine statistical significance versus *S. epidermidis* 1457 wild type (WT), indicated as: not significant (ns), *P < 0.05, **P < 0.01, ***P < 0.001, ****P < 0.0001.

Several other phages, which bind *S. epidermidis* 1457 but cannot replicate in this strain, behaved differently. Siphovirus Φ187, which is only distantly related to ΦE72 and requires GroP-WTA modified with GalNAc for infection of target cells [24], still bound efficiently to the GroP-WTA glucose-deficient mutants (Fig. 5a), indicating that the GroP-WTA glucose modifications are not necessary for Φ187 binding. Φ187 even showed higher plasmid transduction efficiency in the absence of GroP-WTA glucose residues (Fig. 6a). Furthermore, the podoviruses ΦUKE3, ΦSpree, and ΦBE03 [37] exhibited strongly increased binding to the *pgcA, gtaB,* and *tagE* mutants compared to the wild type (Fig. 5 c,f), indicating that these phages are attenuated for binding in the presence of glucose residues on GroP-WTA. Thus, the GroP-WTA glucose residues are important for most of the known *S. epidermidis* phages albeit in quite different ways, depending on the individual phage.

**Figure 6:**
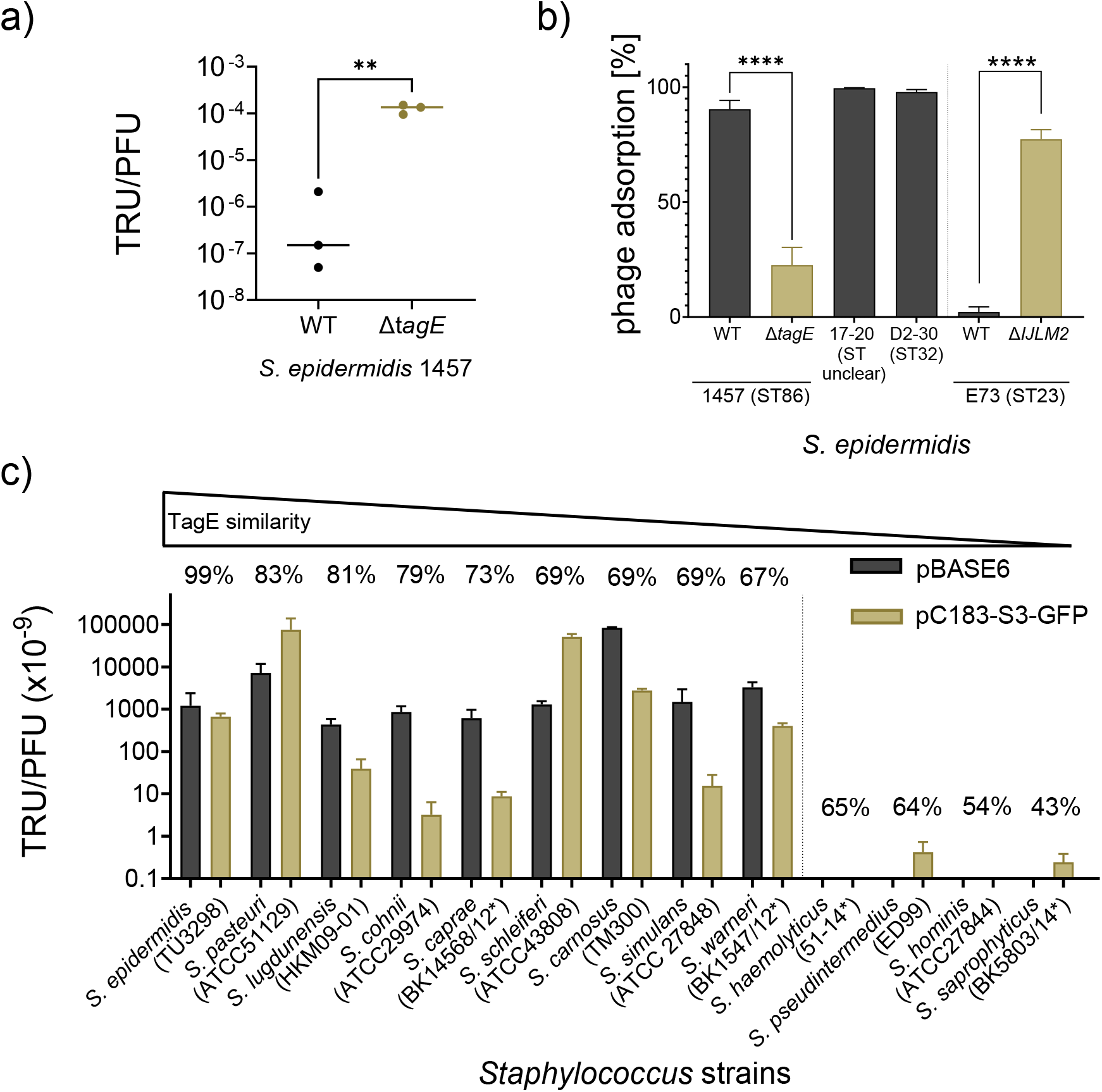
Correlation of TagE-related genes of CoNS species with phage transduction. a) Φ187 transduction of pRB473 is increased in the absence of glucosylated GroP-WTA. b) ΦE72 binds to different strains of *S. epidermidis* but binding is prevented by RboP expression of strain E73. c) ΦE72-mediated transduction of pBASE6 or pC183-S3-GFP to CoNS depends on high TagE homology. If type strains were used to determine sequence similarity of TagE homologues, strain names are marked with an asterisk. The data represents the mean ± SEM of at least three independent experiments. For a,b) unpaired t-test was used to determine statistical significance versus *S. epidermidis* 1457 wild type (WT), indicated as: **P < 0.01, ****P < 0.0001.

### 4. The presence of *tagE* in the genomes of CoNS species corresponds to the capacity of ΦE72 to transduce these species

The *tagE* gene was found in virtually all available *S. epidermidis* genomes suggesting that the substitution of GroP-WTA with glucose is a general trait in *S. epidermidis*. Accordingly, ΦE72 bound well to the tested *S. epidermidis* strains from at least two different sequence types (ST86, ST32) with one exception (Fig. 6b). Notably, ΦE72 did not bind to *S. epidermidis* E73 (ST23), which produces RboP-WTA in addition to GroP-WTA [6]. However, a E73 *tarIJLM2* mutant lacking RboP-WTA was effectively bound by ΦE72 indicating that the additional RboP-WTA shields the surface of *S. epidermidis* in a way that precludes binding of the phage.

GroP-WTA has been reported in several other CoNS species. The nature of the sugar modifications in these species has remained largely unknown, but several CoNS have been reported to contain either glucose, GlcNAc or GalNAc attached to WTA [9]. We succeeded in transducing many different CoNS species via ΦE72 with either the staphylococcal shuttle vector pBASE or the green-fluorescent protein-expressing plasmid pC183-S3 GFP. Some of the available CoNS genomes were found to encode TagE homologs albeit with different degrees of sequence conservation, ranging from 43% to 83% similarity (Table 1). Those species with TagE similarities above 67% could be transduced by ΦE72, while those with less conserved TagE homologs did not take up DNA from ΦE72 (Fig. 6c), suggesting that only CoNS with highly conserved versions of TagE may glycosylate their GroP-WTA in a similar way as in *S. epidermidis* while the others may glycosylate either other WTA backbone types or may transfer other sugars. Among the tested species, *Staphylococcus pasteuri*, *Staphylococcus lugdunensis*, *Staphylococcus cohnii*, *Staphylococcus caprae*, *Staphylococcus schleiferi*, *Staphylococcus carnosus*, *Staphylococcus simulans,* and *Staphylococcus warneri* strains were transducible with ΦE72. Isolates of two of these species*, Staphylococcus cohnii* and *Staphylococcus warneri,* have indeed previously been described to produce GroP-WTA, which is modified with glucose [9]. In contrast to the varying degrees of conservation of *tagE*, the *pgcA* and *gtaB* genes are present in virtually all *Staphylococcus* genomes with high sequence similarity, including *S. aureus*, probably because UDP-glucose is required in all these species for DGlcDAG glycolipid synthesis [36]. Among the strains that encode highly conserved TagE homologues, *tagE* was encoded in the vicinity of both *pgcA* and *gtaB* only in *S. pasteuri* and *S. lugdunensis,* in addition to *S. epidermidis* (Table 1). Thus, phage ΦE72 represents a helpful tool for studying WTA properties and an attractive vehicle for interspecies transduction of DNA among members of the genus *Staphylococcus*.

**Table 1:** Conservation of TagE homologs in CoNS strains used for transduction.

| Species | Strain name | Query cover [%] | Sequence identity [%] | Sequence similarity [%] | *tagE*, *pgcA*, *gtaB* encoded together |
| --- | --- | --- | --- | --- | --- |
| *Staphylococcus epidermidis* | TÜ3298 | 100% | 99% | **99%** | Yes |
| *Staphylococcus epidermidis* | D2-30 | 100% | 99% | **99%** | Yes |
| *Staphylococcus pasteurii* | ATCC51129 | 100% | 69% | **83%** | Yes |
| *Staphylococcus lugdunensis* | HKU09-01 | 99% | 64% | **81%** | Yes |
| *Staphylococcus cohnii* | ATCC29974 | 99% | 61% | **79%** | No |
| *Staphylococcus caprae* | ATCC35538 | 100% | 52% | **73%** | No |
| *Staphylococcus schleiferi* | ATCC43808 | 98% | 50% | **69%** | No |
| *Staphylococcus carnosus* | TM300 | 99% | 48% | **69%** | No |
| *Staphylococcus simulans* | ATCC27848 | 99% | 46% | **69%** | No |
| *Staphylococcus warneri* | ATCC27836 | 99% | 47% | **67%** | No |
| *Staphylococcus haemolyticus* | ATCC29970 | 99% | 45% | **65%** | No |
| *Staphylococcus pseudointermedius* | ED99 | 100% | 43% | **64%** | No |
| *Staphylococcus hominis* | ATCC27844 | 65% | 28% | **54%** | No |
| *Staphylococcus saprophyticus* | ATCC15305 | 85% | 24% | **43%** | No |

## Discussion

WTA structures are known to be highly diverse among Firmicutes, often with species- or even clone-specific composition [7, 38]. Most *S. epidermidis* produce a WTA type that is entirely different from that of *S. aureus* with a GroP rather than a RboP backbone. This study shows that *S. epidermidis* uses a GroP backbone with unmodified or with alanylated glucose. It remains unclear why *S. epidermidis* and *S. aureus* have developed such entirely different WTA types. The different structures may limit the number of bacteriophages that can infect and harm either one or both species. However, ΦK, one the most lytic bacteriophages, can lyse *Staphylococcus* cells irrespective of the WTA backbone structure and a recent study has demonstrated that several *Staphylococcus* phages can infect both, *S. aureus* and *S. epidermidis* [39]. The differences in WTA may limit infections and concomitant lysogenization or transduction events by specific members of the siphovirus group, which depend much more on a specific WTA backbone and glycosylation type than myoviruses. Notably, the presence of glucose on GroP-WTA prevented adsorption to *S. epidermidis* by all tested podoviruses (ΦUKE3, ΦSpree, and ΦBE03). The number of available *S. epidermidis*-targeting phages is still very limited, which impedes more extensive studies on the susceptibility of *S. epidermidis* wild-type and WTA mutant strains for different phage types. Discovery programs for identification of new phages that can infect *S. epidermidis* will help to clarify these questions in the future.

WTA is an important bacterial ligand for host receptors on mammalian immune cells with critical roles in innate immunity [8, 40]. WTA glycosylated with GlcNAc can activate the scavenger receptor langerin on skin Langerhans cells [41]. *S. aureus* is found on the skin of atopic dermatitis patients eliciting skin inflammation in a process that probably involves WTA-langerin interaction [8]. In contrast, *S. epidermidis* cannot activate langerin [41], probably because its GroP-WTA is glycosylated with glucose. It may be advantageous for *S. epidermidis*, one of the most abundant skin-colonizers [1], and for other CoNS, to avoid skin inflammation by producing a non-inflammatory WTA type decorated with glucose.

*S. epidermidis* uses the same pathway for GroP-WTA glycosylation with glucose residues as described for *B. subtilis* [28]. Activation of glucose via the PgcA and GtaB enzymes yields UDP-glucose as donor of glucose residues, which are subsequently transferred to the WTA backbone by TagE. Other WTA glycosyltransferases apart for TarM [27], including those transferring glucose to RboP-WTA in *B. subtilis* W23 (TarQ) [7, 11], GlcNAc to RboP-WTA in certain *S. aureus* clones (TarS, TarP) [11, 42], or GalNAc to GroP-WTA in *S. aureus* ST395 (TagN) [24] share no or very low similarity with TagE. However, protein structure prediction with Alphafold 2 revealed that TagE most likely forms a symmetric, propeller-like homotrimer with each monomer divided into the characteristic glycosyltransferase domain and the β-sheets containing trimerization domain as previously described for the well-studied *S. aureus* glycosyltransferase TarM (Fig. S5) [43–45].

In addition to glucose, WTA is usually also modified with D-alanine [38]. Since GroP repeating units have only one free hydroxyl group for substitution with either D-alanine or glucose, it is not surprising that only ca. 50% of the repeating units carried glucose residues. The teichoic acid D-alanylation machinery attaches D-alanine to a variety of different molecules including LTA, RboP-WTA, and GroP-WTA [46]. Its limited specificity for acceptor substrates may explain why a minor portion of the glucose residues on *S. epidermidis* GroP-WTA are also alanylated. GroP repeating units are shorter than RboP repeating units, which may explain why the additional RboP-WTA polymers of *S. epidermidis* E73 are probably longer and precluded binding of ΦE72 to strain E73. The additional WTA may, therefore, represent a further strategy to interfere with phage infection and with interaction of other WTA-binding molecules.

Several other CoNS species appear to produce a similar WTA type as *S. epidermidis* because they encode potential TagE proteins and interact with ΦE72. Interspecies horizontal gene transfer via WTA-binding transducing phages appears to be rather common among these species and may have contributed to the import of the methicillin-resistance conferring *mecA* gene into *S. epidermidis* and, eventually, to *S. aureus* to create MRSE and MRSA clones [20]. It remains mysterious how the barrier for horizontal gene transfer between *S. epidermidis* and *S. aureus* that results from the substantial differences in WTA structure could be overcome. Specific *S. epidermidis* clonal lineages with both, GroP-WTA and *S. aureus*-type RboP-WTA such as ST10, ST23, and ST87 [6] or the *S. aureus* lineage ST395 producing CoNS-type GroP-WTA [14], may represent critical hubs for the exchange of genetic material between the species *S. epidermidis* and *S. aureus*. Several CoNS species encode potential WTA glycosyltransferase homologs with only low or no similarity to TagE. They may produce other WTA backbones or glycosylate their WTA with other sugars.

*S. epidermidis* often causes difficult-to-treat biofilm-based infections on implanted materials, which frequently require surgical replacement [4]. Treatment with lytic bacteriophages that could destroy *S. epidermidis* biofilms hold promise for the development of new therapeutic strategies [3, 18]. Understanding how phages detect suitable host bacteria and which *S. epidermidis* clones express corresponding phage-binding WTA motives will be important for the success of such strategies. The TagE-mediated WTA glycosylation with glucose might contribute to the narrow host range of lytic podoviruses like ΦBE03 [37]. Accordingly, finding podoviruses, which bind to GroP-WTA glucose might help to develop efficient therapeutic phage cocktails. Moreover, glycosylated WTA is a major antigen for protective antibodies against *S. aureus* [42, 47, 48] and, probably, also *S. epidermidis*. It represents therefore a particularly attractive antigen for vaccine development [48]. As for phage therapy, the success of such vaccination strategies will depend on in-depth knowledge on the structure and prevalence of WTA glycoepitopes among different *S. epidermidis* lineages. Our study may motivate more extensive investigations on WTA glycoepitopes in different staphylococcal pathogens and commensals.

## Materials and Methods

### Bacterial strains and growth conditions

*S. epidermidis* and *S. aureus* strains were cultivated in basic medium (BM) and incubated at 37°C on an orbital shaker. *E. coli* strains were cultivated in lysogeny broth (LB). Media were supplemented with appropriate antibiotics chloramphenicol (10 μg/ml), or ampicillin (100 μg/ml). *E. coli* DC10b and *S. aureus* PS187 Δ*sauUSI*Δ*hsdR* were used as cloning hosts, *S. epidermidis* 1457 was used for gene deletion studies. Bacteriophages and propagation strains used in this study are listed in Table S1.

### Transposon mutagenesis of *S. epidermidis* strain 1457

The transposon plasmid pBTn described previously [25] was used to create a transposon library in *S. epidermidis* 1457. The features of this temperature-sensitive *E. coli/S. aureus* shuttle vector include a mini-transposon with an erythromycin resistance cassette flanked by inverted repeats from the horn fly transposon and a xylose-inducible transposase Himar1, which can mobilize the mini-transposon and integrate it into the chromosome with no bias for any specific sequence. Transposon library construction has been described in detail before [27]. In short, *S. epidermidis* 1457 was transformed with pBTn followed by mobilization of the mini-transposon into the genome upon xylose induction of the transposase. The pBTn plasmid was cured via shifts to nonpermissive temperature.

### Isolation of phage-resistant transposon mutants

To isolate phage-resistant mutants, the transposon mutant library was infected with ΦE72 at a MOI of at least 100. After incubation for up to 4 h, the cells were centrifuged at 5,000 × g for 10 min and plated on TSA agar containing erythromycin. Single colonies of surviving mutants were transferred to fresh TSA agar plates repeatedly. Phage resistance was confirmed by spot assays with ΦE72, and the phage-resistant mutants were treated with 1 µg/ml mitomycin to test for and to exclude lysogeny. To identify the site of transposon insertion, total DNA was isolated, purified with the NucleoSpin® tissue kit (Macherey-Nagel, Düren), digested, religated, multiplied with primers erm-For and erm-Rev (Table S2), which anneal to the erythromycin resistance cassette of the mini-transposon, and sequenced.

### Molecular genetic methods

For the construction of the *tagE*, *pgcA,* and *gtaB* mutants in *S. epidermidis* 1457, the pBASE6 *E. coli/S.aureus* shuttle vector was used according to standard procedures [49]. For mutant complementation, plasmid pRB473 was used [50]. The primers for knockout and complementation plasmid construction are listed in (Table S2). Both pBASE6 and pRB473 containing either the respective up- and downstream fragments for knockout construction (pBASE6) or the complementation sequence (pRB473), were used to transform *E. coli* DC10b, and subsequently PS187 Δ*sauUSI*Δ*hsdR* by electroporation. The plasmids were subsequently transferred to *S. epidermidis* strain 1457 by transduction with Φ187 using *S. aureus* PS187 Δ*sauUSI*Δ*hsdR* as donor strain as described previously [51].

### Phage binding, infection, and transduction assays

Phage spot assays were performed as described previously [14]. All applied bacteriophages (Table S1) were propagated in suitable bacterial host strains and phage lysates were filtered to yield sterile phage suspensions. Test bacteria were cultivated overnight in fresh BM. OD_600_ = 0.1 was adjusted in 5 ml LB soft agar for the preparation of bacterial overlay lawns. 10 µl of phage suspensions were spotted onto the bacterial lawns. After overnight incubation at 37°C for podoviruses and siphoviruses, and 30°C for myoviruses, phage clearing zones and individual plaques were observed and recorded.

Phage adsorption efficiency was determined as described previously with minor modifications [14]. Briefly, adsorption rates were analyzed by mixing approximately 10^6^ PFU/ml in BM supplemented with 4 mM CaCl_2_ with the tested bacteria at an OD_600_ of 0.5 and incubating for 15 min at 37°C. The samples were subsequently centrifuged, and the supernatants were spotted on indicator strains to determine the number of unbound phages in the supernatant. The adsorption rate was calculated by dividing the number of bound phages by the number of input phages.

Transduction experiments were performed as described previously [14]. Briefly, 1 ml of exponentially growing cultures of a recipient strain was adjusted to an OD_600_ of 0.5. The cells were sedimented by centrifugation and resuspended in 200 µl of phage buffer containing 0.1% gelatin, 1 mM MgSO_4_, 4 mM CaCl_2,_ 50 mM Tris, and 0.1 M NaCl. 200 µl of bacteria in phage buffer were mixed with 100 µl of lysates obtained from *S. aureus* PS187 and *S. epidermidis* 1457 donor strains carrying plasmids of choice. Samples were then incubated for 15 min at 37°C, diluted, and plated on chloramphenicol-containing BM agar to count colonies.

### Electron microscopy

*S. epidermidis* 1457 wild type, *ΔtagE*, *ΔpgcA*, and *ΔgtaB* were grown until stationary phase, and fixed at an OD_600_ of 10 in 200 µl Karnovsky’s fixative (3% formaldehyde, 2.5% glutaraldehyde in 0.1 M phosphate buffer pH 7.4) for 24 h. Samples were then centrifuged at 1,400 x g for 5 min, supernatant was discarded, pellets were resuspended in approximately 20 µl agarose (3.9%) at 37°C, cooled to room temperature, and cut into small pieces. Postfixation was based on 1.0% osmium tetroxide containing 2.5% potassium ferrocyanide (Morphisto) for 2 h. After following the standard methods, samples were embedded in glycide ether and cut using an ultramicrotome (Ultracut E, Reichert). Ultra-thin sections (30 nm) were mounted on copper grids and analyzed using a Zeiss LIBRA 120 transmission electron microscope (Carl Zeiss) operating at 120 kV.

### WTA isolation

WTA was isolated as described previously [14, 52, 53] with minor modifications. Briefly, bacterial cells from two liters of overnight cultures were washed and disrupted with glass beads in a cell disrupter (Euler). Cell lysates were incubated at 37°C overnight in the presence of DNase and RNase. SDS was added to a final concentration of 2% followed by ultrasonication for 15 min. Cell walls were washed several times to remove SDS. To release WTA from cell walls, samples were treated with 5% trichloroacetic acid for 4 h at 65°C. Peptidoglycan debris was separated via centrifugation (10 min, 14,500 x g). Determination of phosphate amounts as described previously [53–55] was used for WTA quantification. Crude WTA extracts were further purified as already described [27]. Briefly, the pH of the crude extract was adjusted to 5 with NaOH and dialyzed against water with a Slide-A-Lyzer Dialysis Cassette (MWCO of 3.5 kDa; Thermo Fisher Scientific). For HPLC-MS analysis, 50 µl of 100 mM WTA sample were hydrolyzed with 100 mM NaOH at 60°C for 2 h. The remaining dialyzed WTA was further lyophilized for long-term storage at −20 °C or used for further analysis. 10-15 mg lyophilized WTA sample were used for NMR. Detailed explanations of the HPLC-MS and NMR methods can be found as extended descriptions of detailed methods.

### Enzymatic determination of glucose in the WTA samples

The High Sensitivity Glucose Assay Kit (mak181, Sigma-Aldrich) was used to determine the glucose content in the WTA sample. 50 µl of dialyzed WTA samples and 50 µl of 1 mM glucose standard solution were dried in a vacuum concentrator at 60°C. 100 µl of 0.5 M HCl was added to the samples and the standard solution and cooked for 2 h in a water bath. The glucose standard was diluted 1:50 resulting in a 10 µM concentration and different volumes were used to cover a range of 0 - 100 pmol. Samples were also diluted at least 1:50 and different dilutions of the samples were tested in a 96-well plate. The assay was performed according to the manufacturer’s instructions. The fluorescence intensity was measured at excitation wavelength 535 nm and emission wavelength 587 nm.

### Glycolipid isolation, thin layer chromatography (TLC) and detection with α-naphthol

The detection of glycolipids was performed similar to a previously described method [36]. *S. epidermidis* 1457 and the respective mutants were grown to OD_600_ of 3.5. 5 ml of bacterial suspension were washed and resuspended in 500 µl of 100 mM sodiumacetate (pH 4.7) and transferred into glass vials. 500 µl chloroform and 500 ml methanol were added and the mixture was vortexed vigorously. The samples were centrifuged at 4,600 x g for 20 min at 4°C and the lower phase was dried overnight and resuspended in 25 µl methanol and chloroform in a 1:1 ratio. The whole sample was applied to a high-performance thin-layer chromatography (HPTLC) silica gel 60 plate (10 x 10 cm; Merck) with a Hamilton syringe. A positive control containing 5 µg digalactosyldiacylglycerol (DGDG, Sigma-Aldrich) was used. A Linomat 5 (Camag), and an auto developing chamber (Camag), were used to apply the sample to the TLC plate and to run it with a solvent containing 65:25:4 (v/v/v) chloroform/methanol/H_2_O. The dried TLC plate was sprayed with 3.2% α-naphthol in methanol/H_2_SO_4_/H_2_O 25:3:2 (v/v/v) and the glycolipids were visualized by heating the plate at 110°C for a few minutes.

### In silico analysis

All statistical analyses were performed with Graph Pad Prism 9.2.0 (GraphPad Software, La Jolla, USA). Multiple sequence alignment was performed with SnapGene® 5.3.2 using MUSCLE. Protein structure prediction was done using AlphaFold2 with ColabFold [44, 45].

## Acknowledgements

We thank David Gerlach, Xin Du, and Bernhard Krismer for helpful discussions, Arnaud Kengmo Tchoupa and Ulrike Redel for help with TLC, and Y. Que and E. Baumgartner for supply of phages. This work was financed by grants from the German Research Foundation to A.P. (TRR34; TRR165 project ID 246807620; PE 805/7-1 project ID 410190180; PE 805/8-1 project ID 410190180) and the German Center for Infection Research (DZIF) to A.P. The authors acknowledge infrastructural support by the Cluster of Excellence EXC 2124 ‘Controlling Microbes to Fight Infections’ project ID 39083813

---

**Fig. S1:**
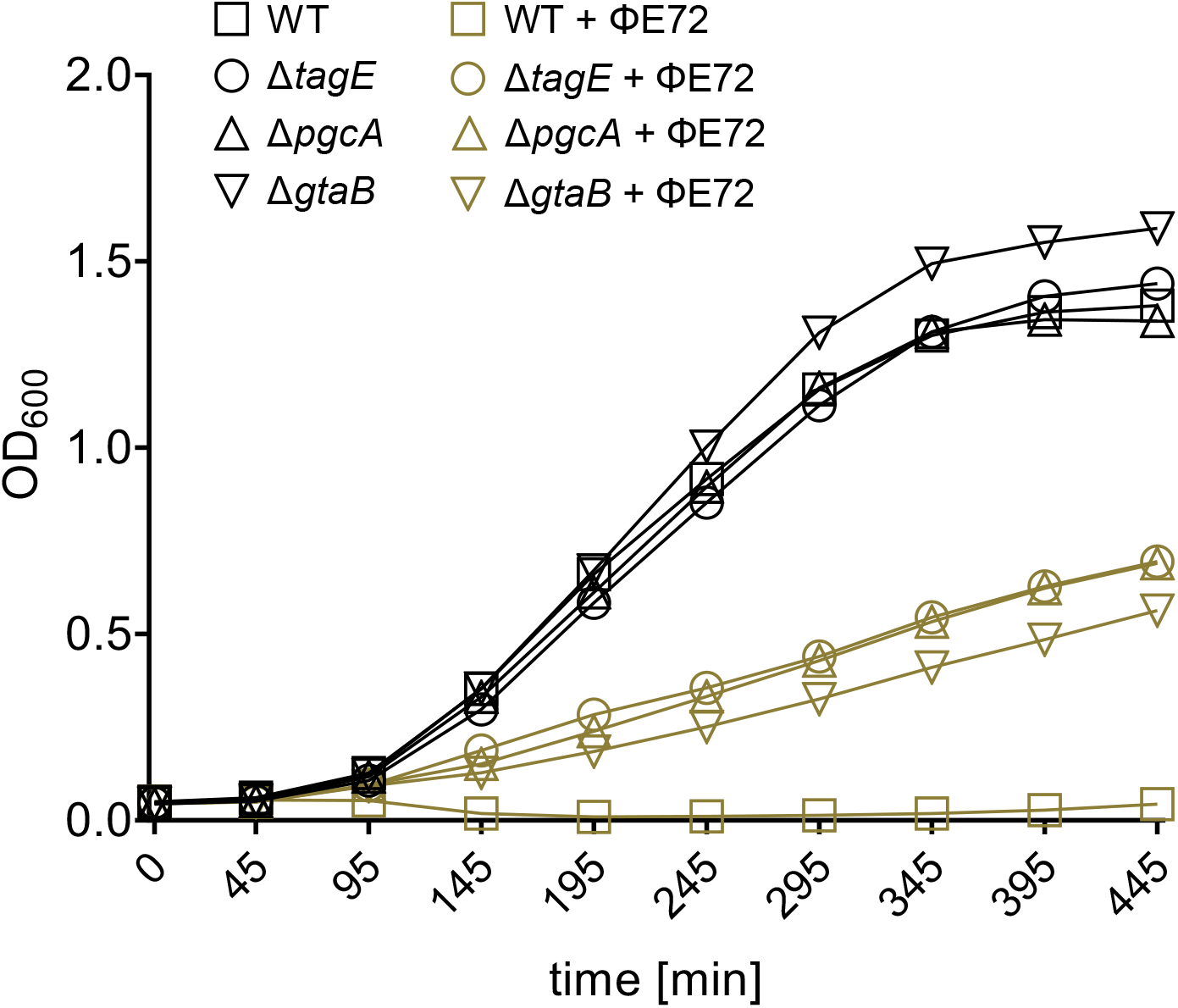
ΦE72 prevents growth of *S. epidermidis* 1457 wild type (WT). Growth of the Δ*tagE*, Δ*pgcA,* Δ*gtaB* mutants is only partially reduced by ΦE72 compared to growth without addition of phage. Approximately 5×10^8^ PFU/ml were used. Data represent mean ± SEM of three independent experiments.

**Fig. S2:**
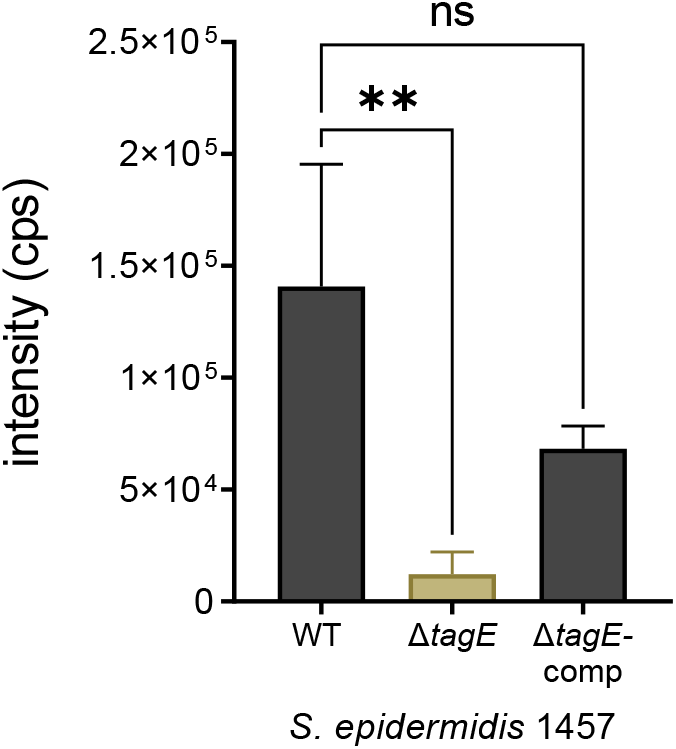
Area-under-the-curve quantification of GroP-GroP-Glc residue ([M - H]^−^ = 487.0623) total ion current (TIC) chromatogram measured by HPLC-MS after chemical digest of *S. epidermidis* WTA. Data represent mean ± SEM of three independent experiments. Ordinary one-way ANOVA was used to determine statistical significance versus *S. epidermidis* 1457 wild type (WT), indicated as: not significant (ns), **P < 0.01.

**Fig. S3:**
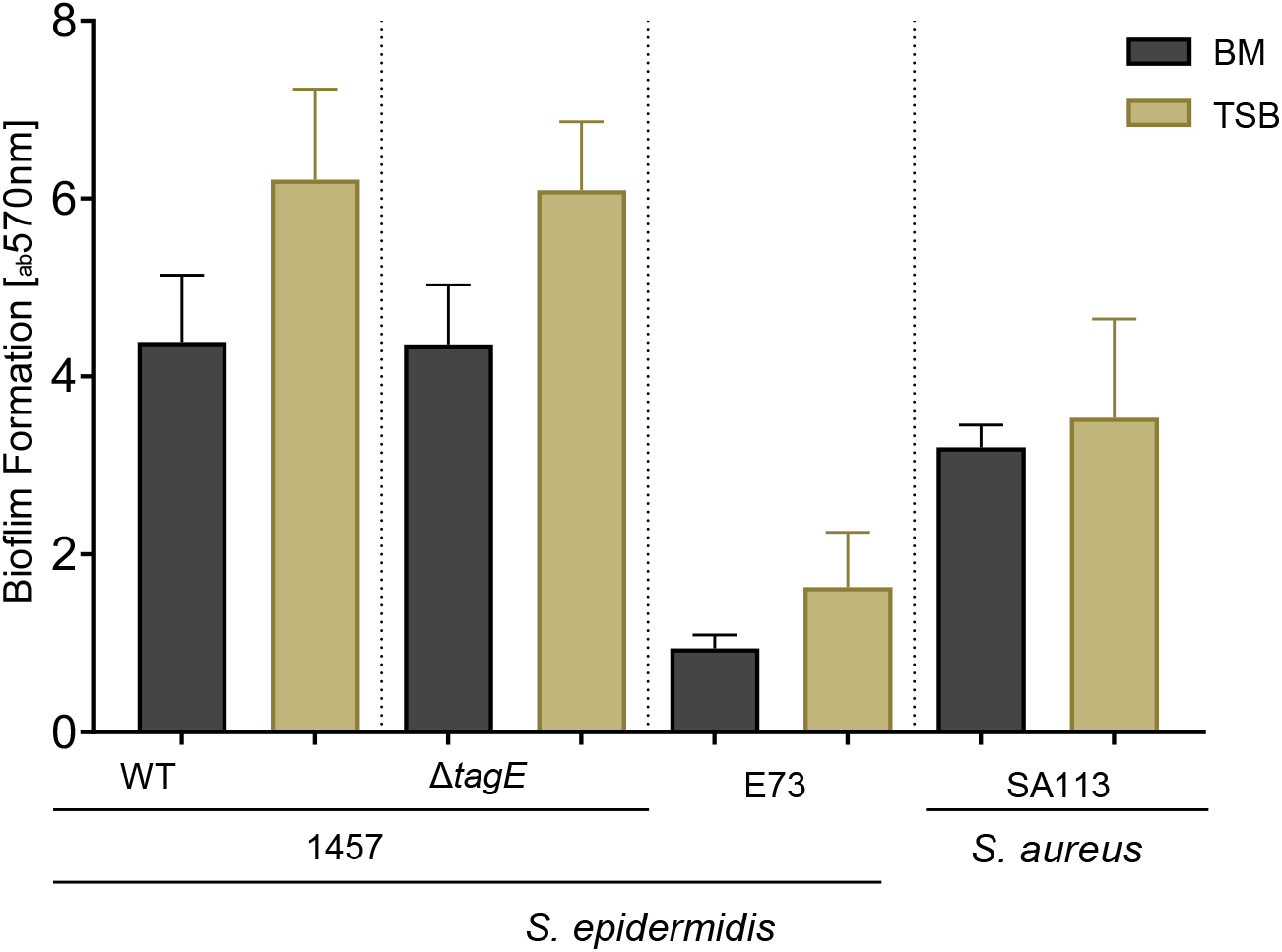
*S. epidermidis* 1457 biofilm formation was measured in BM and TSB medium. Biofilm formation is unchanged in the Δ*tagE* deletion mutant.

**Fig. S4:**
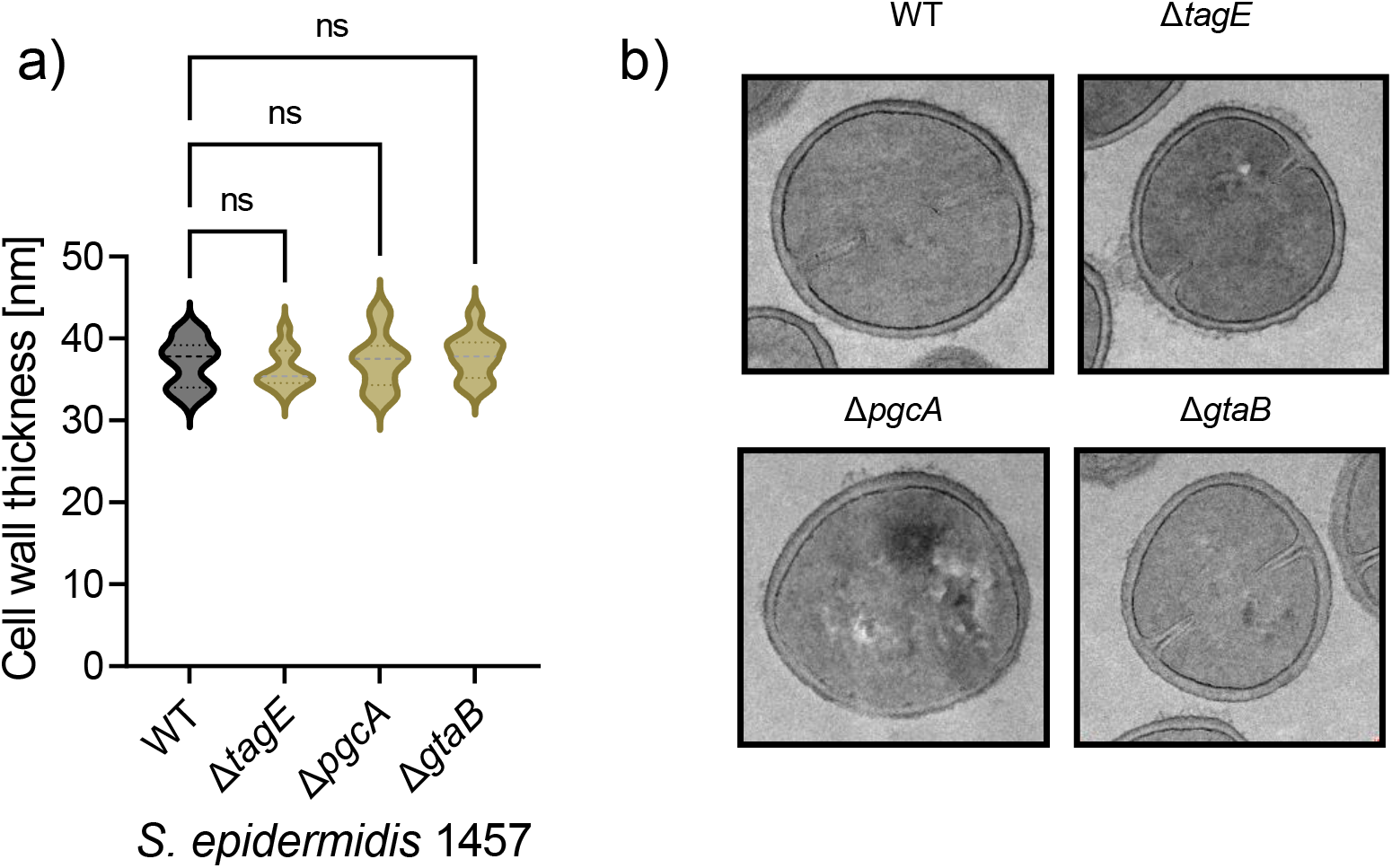
Electron microscopy at 12,500 x magnification indicates that cell wall thickness (a), and cell shape (b), is unchanged in all mutants compared to the wild type. a) shows the mean cell wall thickness of at least 11 different bacterial cells of each mutant or the wild type (WT). Ordinary one-way ANOVA was used to determine statistical significance versus *S. epidermidis* 1457 wild type (WT), indicated as: not significant (ns).

**Fig. S5:**
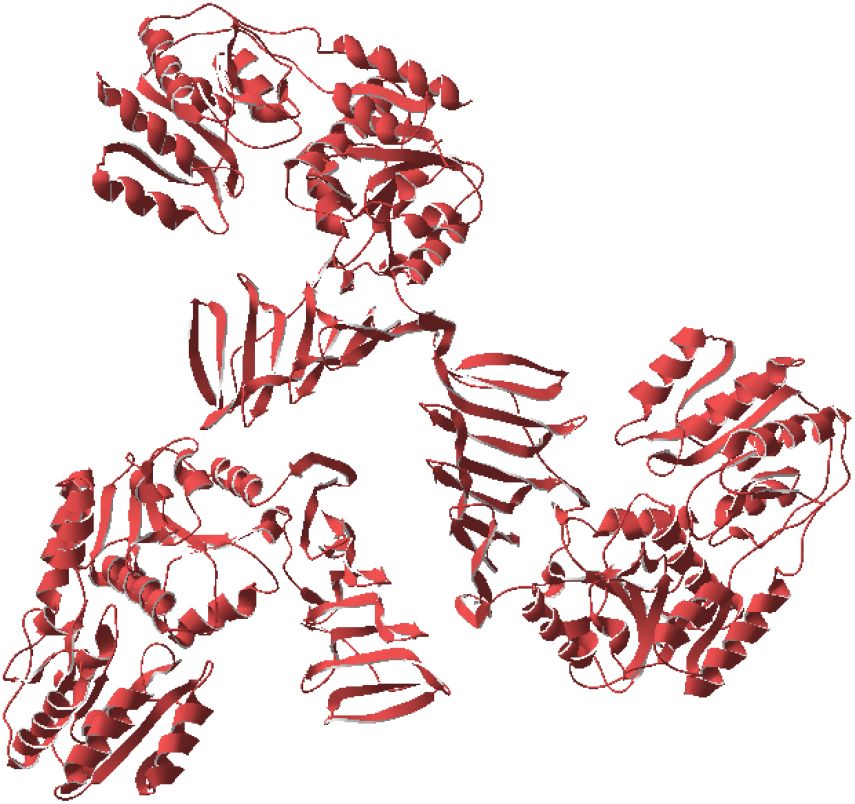
Structural prediction of the *S. epidermidis* TagE trimer with Alphafold2 [44, 45].

**Table S1:**
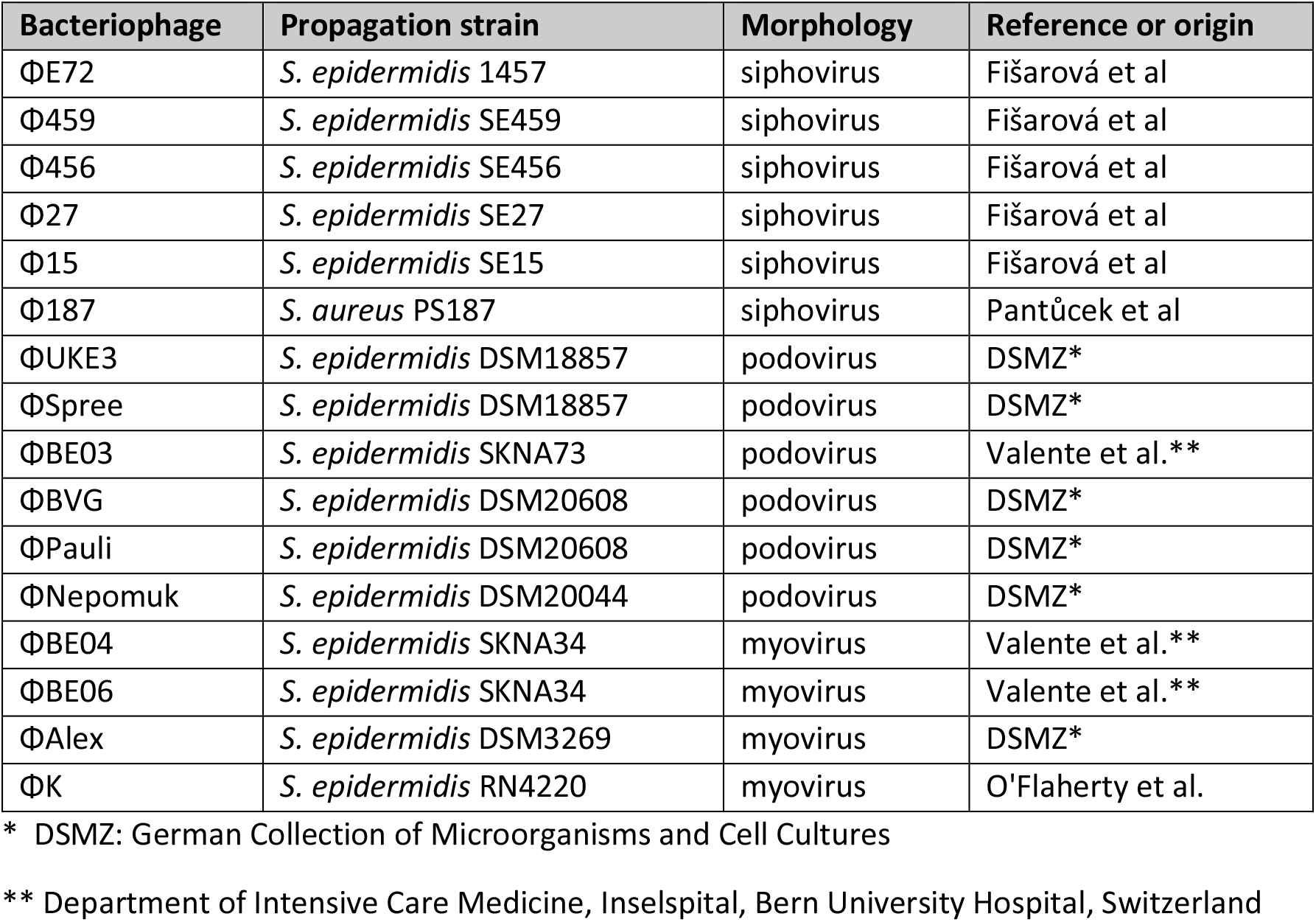
Bacteriophages and bacterial strains used in this study.

| Bacteriophage | Propagation strain | Morphology | Reference or origin |
| --- | --- | --- | --- |
| ΦE72 | *S. epidermidis* 1457 | siphovirus | Fišarová et al |
| Φ459 | *S. epidermidis* SE459 | siphovirus | Fišarová et al |
| Φ456 | *S. epidermidis* SE456 | siphovirus | Fišarová et al |
| Φ27 | *S. epidermidis* SE27 | siphovirus | Fišarová et al |
| Φ15 | *S. epidermidis* SE15 | siphovirus | Fišarová et al |
| Φ187 | *S. aureus* PS187 | siphovirus | Pantůček et al |
| ΦUKE3 | *S. epidermidis* DSM18857 | podovirus | DSMZ* |
| ΦSpree | *S. epidermidis* DSM18857 | podovirus | DSMZ* |
| ΦBE03 | *S. epidermidis* SKNA73 | podovirus | Valente et al.** |
| ΦBVG | *S. epidermidis* DSM20608 | podovirus | DSMZ* |
| ΦPauli | *S. epidermidis* DSM20608 | podovirus | DSMZ* |
| ΦNepomuk | *S. epidermidis* DSM20044 | podovirus | DSMZ* |
| ΦBE04 | *S. epidermidis* SKNA34 | myovirus | Valente et al.** |
| ΦBE06 | *S. epidermidis* SKNA34 | myovirus | Valente et al.** |
| ΦAlex | *S. epidermidis* DSM3269 | myovirus | DSMZ* |
| ΦK | *S. epidermidis* RN4220 | myovirus | O'Flaherty et al. |

\* DSMZ: German Collection of Microorganisms and Cell Cultures
\*\* Department of Intensive Care Medicine, Inselspital, Bern University Hospital, Switzerland

**Table S2:**
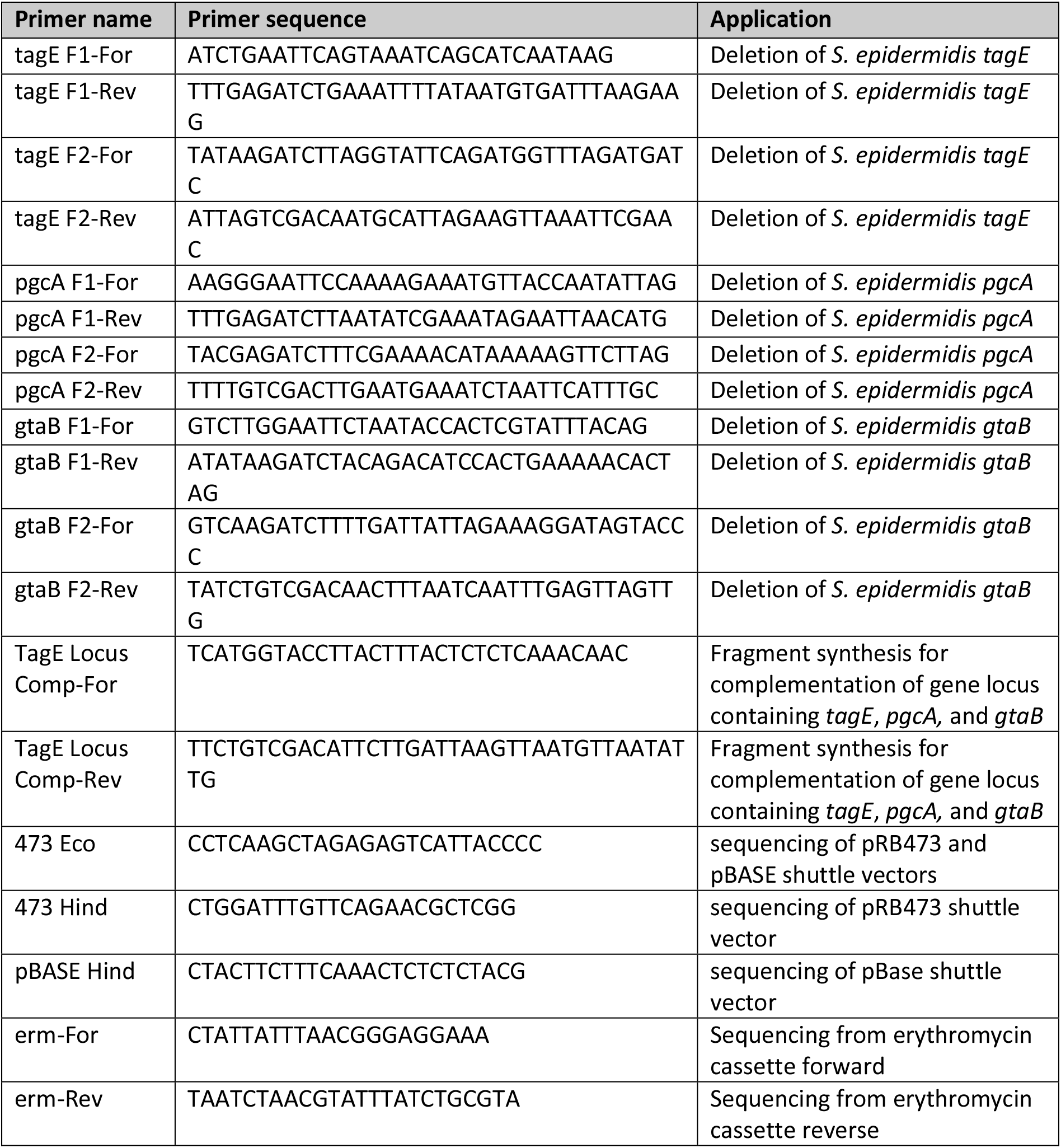
Primer sequences used for cloning and sequencing.

## Extended descriptions of detailed methods

### WTA compositional analysis

#### HPLC-MS

Analysis of the WTA polymer composition was performed using an LTQ Orbitrap Velos mass spectrometer (Thermo Fisher Scientific), connected to an ACQUITY ultra-performance liquid chromatography (UPLC) system (Waters Corporation). Separation in the UPLC was carried out using a Phenomenex C18-Gemini® column (150 × 2 mm, 3 μm, 110 Å, Phenomenex) at 37°C with 0.1% formic acid and 0.05% HCO_2_NH_4_ (A) and CH_3_CN (B) buffer system. A single run (injection volume of 5 μl) was performed with a flow rate of 0.2 ml/min and a two-step gradient: after 2.5 min of equilibration with 100% A, a 1-min gradient up to 5% B was followed by a 4-min gradient up to 70% B. After 2 min at 70% B, a re-equilibration step of 2.5 min followed with a flow rate of 4 ml/min. LC-MS data processing was done with UmetaFlow GUI ([56], https://github.com/axelwalter/streamlit-metabolomics-statistics) via extracted ion chromatograms with a mass tolerance of 10 ppm.

#### NMR

1H NMR spectra were recorded for both, wild-type and Δ*tagE S. epidermidis* strains, and they were carried out on a Bruker DRX-600 spectrometer equipped with a cryo-probe, at 298 K. Chemical shifts of spectra recorded in D_2_O were calculated in ppm relative to internal acetone (2.225 and 31.45 ppm). 2D NMR spectra were acquired for *S. epidermidis* wild type only, the spectral width was set to 12 ppm and the frequency carrier placed at the residual HOD peak, suppressed by pre-saturation. Two-dimensional spectra (DQ-COSY, TOCSY, NOESY, gHSQC, and gHMBC) were measured using standard Bruker software. For all experiments, 512 FIDs of 2,048 complex data points were collected, 32 scans per FID were acquired for homonuclear spectra, and 100 and 200 ms of mixing time was used for the TOCSY and NOESY spectra, respectively. Heteronuclear ^1^H-^13^C spectra were measured in the ^1^H-detected mode, gHSQC spectrum was acquired with 40 scans per FID, the GARP sequence was used for ^13^C decoupling during acquisition; gHMBC scans doubled those of gHSQC spectrum. During processing, each data matrix was zero-filled in both dimensions to give a matrix of 4K × 2K points and was resolution-enhanced in both dimensions by a cosine-bell function before Fourier transformation; data processing and analysis were performed with the Bruker Topspin 3 program.

### NMR analysis of the WTA of the wild type (WT) strain of *Staphylococcus epidermidis*

NMR analyses of the spectra displayed several signals in the anomeric region (5.5 – 4.4 ppm, Fig. 3c) of the proton spectrum with the one at 5.20 ppm being more intense than the others. Then, inspection of the HSQC spectrum (Fig S6a) disclosed that only the signals at ∼ 5.2 and ∼ 5.1 ppm arose by the anomeric position of different monosaccharide residues, due to the characteristic values of the related carbon atoms (Table S3, [57]). The full assignment of both proton and carbon chemical shifts was possible with confidence only for the most abundant unit, labelled with **A.** Thus, the anomeric proton at 5.2 ppm was labelled **A_1_**, and the combined analysis of the TOCSY and COSY spectra determined that it was an α-glucose (Fig. S6b). Indeed, the TOCSY spectrum showed that **A_1_** correlated to four other protons as occurs for *gluco* configured residues, and this information combined with those from the COSY spectrum enabled the sequence assignment from H-2 to H-5 (Fig. S6b, Table S3). Then, the identification of A6 was inferred by the finding of the H-4/H-6 cross peak in the TOCSY spectrum (Fig. S6b) while the position of the other H-6 proton, labelled A6’ was determined by the strong cross-peak in the COSY spectrum (Fig. S6b). Finally, the identification of the carbon chemical shifts was inferred by analysing the ^1^H-^13^C HSQC (Fig. S6a), which determined that **A** was a glucose unit that was not further substituted due to the similarity of its carbon chemical shifts to those reported for the reference glycoside [57]. The inspection of the HMBC spectrum (not shown) reported a cross peak connecting H-1 of **A** to a carbon at 76.7 ppm in turn correlated to a proton at 4.12 ppm, later assigned to H-2/C-2 of a glycerol (Gro) unit, labelled **b**.

Interestingly, H-1 of **A** was flanked by a second anomeric proton at 5.22 ppm (Fig. S6b, Table S3), labelled as **A’** and presenting a correlation pattern in the TOCSY spectrum very similar to that of **A**, except for the fact that the density analogue to **A_1,5_** was missing while there was a new one relating H-1 to a proton at 4.21 ppm. The identification of the sequence between the protons of this second spin system was aided by the COSY spectrum and the additional signal at 4.21 was assigned to H-5, in turn correlated to the two H-6 protons at 4.66 and 4.44 ppm (Fig. S6b), highly deshielded due to the O-acylation with an Ala residue as inferred by the long range correlation with a carbonyl group at 171.5 ppm (not shown).

Then, the anomeric region reported a proton signal at 5.39 ppm, attached to a carbon at 75.5 ppm with only one additional correlation in the COSY spectrum with a proton at ca. 4.1 ppm, assigned with a hydroxy-methyl carbon at 64.9 ppm in the HSQC spectrum (Figure S6a). The pattern of this unit, labelled **a**, was found to be consistent with that of a Gro unit, phosphorylated at both ends and acylated with an Ala unit at O-2, as described in the WTA polymers containing GroP motifs [42].

Finally, the HSQC spectrum contained three densities at ^1^H/^13^C 4.04/70.8, and 4.12/76.7, labelled as **c_2_**, and **b_2_**, respectively, all identified with the aid of the values reported in literature (Table S3). In detail, **c** was a glycerol unit not further substituted [42], while **b** had the glucose units (**A** and **A’**) linked to O-2 [58]. Of note, the HSQC spectrum contained other densities not related to the WTA polymer and presumably belonging to other compounds co-purified with it. In some cases, it was possible to recognize some amino acids, but it was never possible to establish the nature of the compound(s) due to the low intensities of the signals or to the lack of the proper correlations in the full set of NMR spectra acquired. The integration of the **A_5,1_** and **A’_5,1_** densities in the TOCSY spectrum (Figure S6b) revealed that about 15% of this monosaccharide was derivatized with an alanine at O-6.

**Table S3:** NMR chemical shifts. ^1^H (600MHz) chemical shifts of WTA structural motifs found in *S. epidermidis* wild type. The sample was dissolved in deuterated water (HOD, 550 μl) and measured at 298 K. By convention, C-1 of the glycerol unit is placed at the left of the structural formula, P stands for phosphate.

| Residue | motif | 1;1' (for Gro) | 2 | 3; 3' (for Gro) | 4 | 5 | 6; 6' |
| --- | --- | --- | --- | --- | --- | --- | --- |
| **a** |  | 4.11 x 2 | 5.39 | 4.11 x 2 | -- | -- | -- |
| **Gro** | Ala | 64.9 | 75.5 | 64.9 | -- | -- | -- |
| **b** |  | ~ 4.02 x 2* | 4.12 | ~ 4.05 – 4.00* | -- | -- | -- |
| **Gro** | α-Glc (A or A') | 66.6 | 76.7 | 65.8 | -- | -- | -- |
| **c** |  | 3.85;3.90 | 4.04 | 3.85;3.90 | -- | -- | -- |
| **Gro** |  | 67.5 | 70.8 | 67.5 | -- | -- | -- |
| **A** |  | 5.20 | 3.54 | 3.78 | 3.41 | 3.95 | 3.89; 3.77 |
| **t-α-Glc** |  | 98.9 | 72.8 | 74.3 | 71.0 | 73.1 | 61.9 |
| **A'** |  | 5.22 | 3.57 | 3.80 | 3.43 | 4.21 | 4.66; 4.44 |
| **t-α-Glc6Ala** |  | 96.2 | 72.6 | 74.1 | 71.2 | 71.0 | 66.5 |
| **Ala** |  | -- | 4.23 | 1.62 | -- | -- | -- |
|  |  | 171.5 | 50.1 | 16.6-- | -- | -- | -- |

\* These signals can be exchanged.

**Fig. S6:**
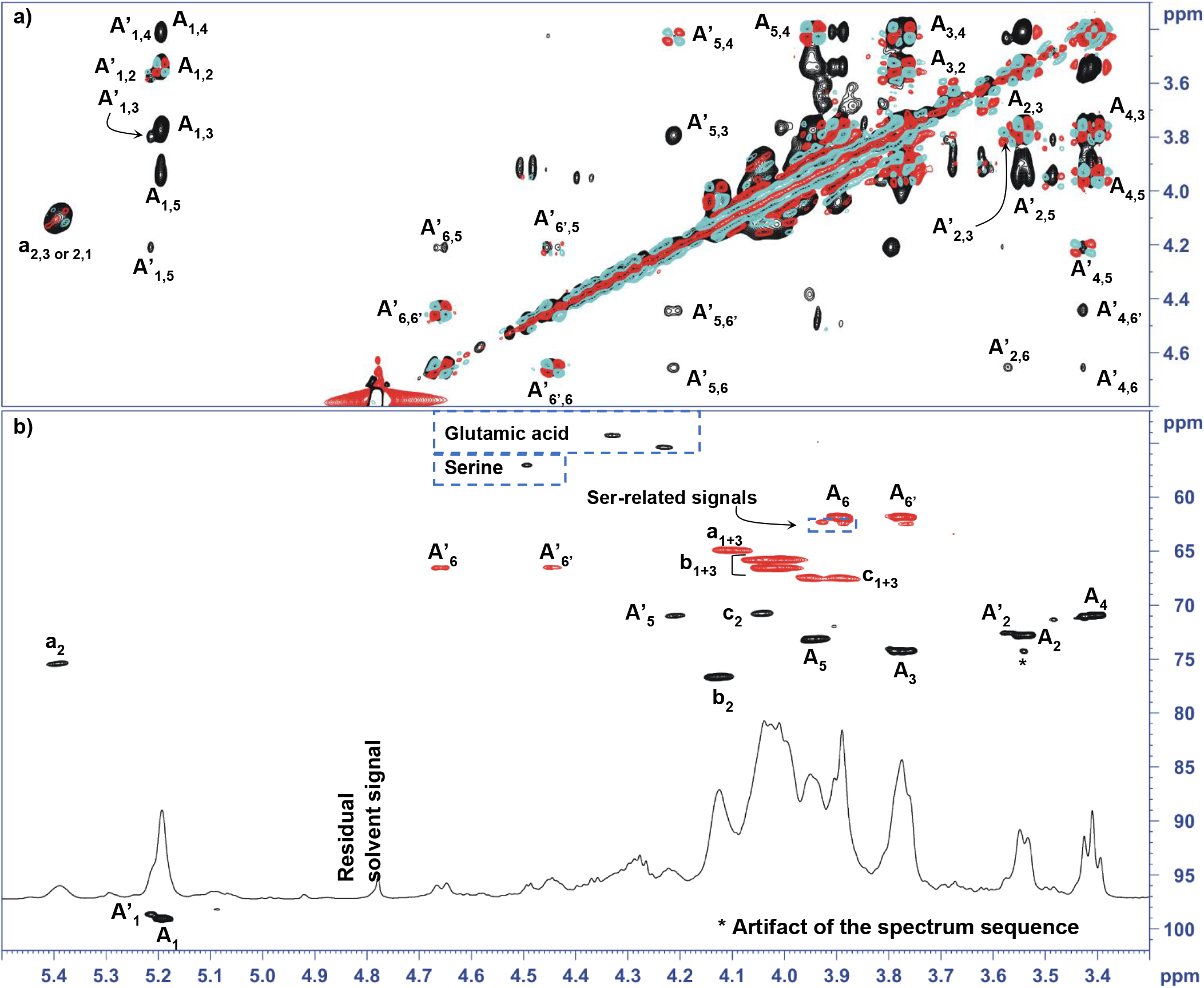
NMR spectra recorded for WTA isolated from *S. epidermidis* wild type. a) Expansion of the HSQC spectrum detailing the anomeric and the carbinolic region. b) Overlap of the TOCSY (black) and COSY (cyan and red) spectra. In all the spectra, the most relevant densities are labelled with the letter used in Table S3; as for the carbohydrate units (**A** and **A’**), the anomeric signals are indicated with a capital letter, while the Gro units (**a**, **b**, and **c**) are labeled with small letters.

